# New cellular models of undifferentiated pleomorphic sarcoma and malignant peripheral nerve sheath tumor

**DOI:** 10.1101/2021.05.13.443902

**Authors:** Angela D. Bhalla, Sharon M. Landers, Anand K. Singh, Michelle G. Yeagley, Gabryella S.B. Meyerson, Zachary A. Mulder, Cristian B. Delgado-Baez, Stephanie Dunnand, Veena Kochat, Katarzyna J. Tomczak, Theresa Nguyen, Xiaoyan Ma, Svetlana Bolshakov, Brian A. Menegaz, Salah-Eddine Lamhamedi-Cherradi, Joseph A. Ludwig, Hannah C. Beird, Xizeng Mao, Xingzhi Song, Davis R. Ingram, Wei-Lien Wang, Alexander J. Lazar, Ian E. McCutcheon, John M. Slopis, Kunal Rai, Jianhua Zhang, Dina C. Lev, Keila E. Torres

## Abstract

Undifferentiated pleomorphic sarcoma (UPS) and malignant peripheral nerve sheath tumor (MPNST) are aggressive soft tissue sarcomas that do not respond well to current treatment modalities. The limited availability of UPS and MPNST cell lines makes it challenging to identify potential therapeutic targets in a laboratory setting. Understanding the urgent need for improved treatments for these tumors and the limited cellular models led us to generate additional cell lines to study these rare cancers further. Patient-derived tumors were used to establish 5 new UPS models, including one radiation-associated UPS—UPS060.1, UPS271.1, UPS511, UPS0103, and RIS620—and 3 new models of MPNST—MPNST007, MPNST3813E, and MPNST4970. This study examined the utility of the new cell lines as sarcoma models by assessing tumorigenic potential and mutation status for known sarcoma-related genes. All the cell lines formed colonies and migrated *in vitro*. The *in vivo* tumorigenic potential of each cell line was determined by either subcutaneous injection of cells or implantation of tumor tissue into immunocompromised mice. UPS060.1, UPS271.1, and UPS511 cells formed tumors in mice upon subcutaneous injection. UPS0103 and RIS620 tumor implants formed tumors *in vivo*, as did MPNST007 and MPNST3813E tumor implants. Mutation analysis of a panel of genes frequently mutated in sarcomas showed that two of the three MPNST cell lines had *NF1* mutations. Two of the three MPNST cell lines had mutations in polycomb repressive complex 2 members. These new cellular models provide the scientific community with powerful tools for detailed studies of sarcomagenesis and investigate novel therapies for UPS and MPNST.

---

Soft tissue sarcomas are rare malignancies originating from the mesoderm. There exist more than 50 histological subtypes based on their tissues of origin. Undifferentiated pleomorphic sarcoma (UPS) comprises a subset of 5% to 15% of soft tissue sarcomas for which clinicians cannot determine the tissue of origin from analysis of the tumor’s molecular and histologic characteristics ^1 2 3^. The standard of treatment for UPS is surgical resection. For patients in whom surgery is not feasible, chemotherapy and radiation therapy may be used, but they have low-to-moderate response rates ^4, 5^. UPS is characterized by a high level of genomic instability, as indicated by its complex karyotype. However, specific recurring genetic alterations are uncommon ^6^. Malignant peripheral nerve sheath tumors (MPNSTs), which comprise 2% of all soft tissue sarcomas ^1^, are another rare and genomically unstable sarcoma. Half of MPNSTs occur in patients with the autosomal dominant cancer predisposition syndrome neurofibromatosis type 1 (NF1; OMIM # 162200). Approximately 40% of MPNST cases are sporadic, and the remaining 10% are associated with previous radiation therapy ^7–9^. Chemotherapy and radiation therapy are ineffective for treating MPNSTs ^2, 10, 11^; therefore, surgical resection remains the standard treatment ^12–15^. MPNSTs have a high rate of local recurrence ^16, 17^. The 5-year survival rate is 35% to 50%, and the 10-year survival rate is 7.5%, stressing the need for improved treatment options ^9, 18^. Limited treatment options are available for patients with UPS and MPNST, largely because the mechanisms involved in the progression of these heterogeneous sarcomas require further investigation ^9, 19, 20 21^.

UPS and MPNST both have complex karyotypes, and no specific genomic alterations are unique to the tumor type. UPS has complex cytogenetic rearrangements in 30% to 35% of the genome ^22^, but to date, no specific alterations have been useful for classifying UPS. Loss of chromosome 13q, which leads to inactivation of the RB pathway, is frequently observed in UPS ^23–25^, and losses of genes in the TP53/ARF/MDM2 pathway are also common ^26, 27^. A study of soft tissue sarcoma by The Cancer Genome Atlas Research Network ^28^ reported several somatic copy number alterations, specifically deep deletions in *RB1* in 16%, *TP53* in 16%, and *CDKN2A* in 20% of UPS. The mutation burden was low in soft tissue sarcomas, an average of 1.06 per MB in 206 tumors. The significantly mutated genes in UPS included *RB1* and potential driver truncating mutations of *NF1* ^28^. MPNST’s complex genomic profile includes copy number aberrations, like microdeletions of *NF1* in 5% to 20% of cases ^29, 30^, as well as *CDKN2A* inactivation ^31, 32^, and *TP53* mutations. Recently, mutations in the genes of the polycomb repressive complex 2 (PRC2) components *SUZ12* and *EED* and the loss of histone 3 lysine 27 trimethylation (H3K27me3) were proposed as markers for NF1-related and sporadic MPNSTs ^20, 33^.

Cell line models that can form tumors in mice are valuable tools to advance the understanding of soft tissue sarcomas with no known genetic drivers such as UPS and MPNST. Currently, some UPS and radiation-induced sarcoma (RIS) cell lines are available for study of these tumor types: U2197, NCC-UPS2-C1 ^34, 35^, 2 developed by our laboratory, UPS186 and RIS819 ^21^, and others, some with the ability to form tumors when injected subcutaneously into mice ^36, 37^. Well-studied NF1-associated MPNST cell lines are also available within the scientific community, including SNF02.2 and SNF94.3, both derived from a metastatic site in the lung, SNF96.2, isolated from a recurrent MPNST; S462, ST88-14, T265, and MPNST642 ^38, 39 40, 41^. All of the NF1-associated MPNST cell lines listed above can form tumors in mice, allowing for the *in vivo* study of these difficult-to-treat sarcomas ^38, 39, 42–44^. However, further development of UPS and MPNST cell line models is warranted because cell lines from genomically complex sarcomas are underrepresented in cancer cell line databases such as the Cancer Cell Line Encyclopedia (CCLE) and Catalogue Of Somatic Mutations In Cancer COSMIC ^45, 46^.

This study aimed to generate and characterize new cellular models of UPS and MPNST that will deepen our understanding of the aberrant cellular processes of these sarcomas known to lack clear genetic or molecular markers. The UPS and MPNST cell lines generated in this study will be available to the scientific community to facilitate the study of these debilitating tumors.

## MATERIALS AND METHODS

### UPS and MPNST primary culture and cell lines

We obtained approval from the Institutional Review Board of The University of Texas MD Anderson Cancer Center and patients’ informed written consent before establishing the cell lines described in this study. The cell lines were isolated from surgical specimens from patients at MD Anderson Cancer Center. A sarcoma pathologist confirmed the final histology of the tumors at MD Anderson Cancer Center. A portion of the original specimens was fixed in formalin and paraffin-embedded, followed by immunohistochemical (IHC) staining for hematoxylin and eosin to visualize the samples. Isolation of tumor cells for RIS620, UPS271.1, and UPS511 was described previously by May et al. UPS060.1 and UPS0103 cells were isolated as described by Peng et al. ^47^. Briefly, fresh sterile samples from resected tumors were minced in Dulbecco’s modified Eagle medium (DMEM; Corning, cat # 10013CV and incubated with collagenase type I (3%; Sigma-Aldrich, cat # C-0130), DNase I (0.02%; Sigma-Aldrich, cat # D-4527), and hyaluronidase (1.5 mg/ml; Sigma-Aldrich, cat # H-3506) at 37°C for 2 to 4 h. After digestion, the sample was strained through a mesh screen (BD Biosciences, cat # 352360), and undigested tissue was discarded. The sample was centrifuged, washed, resuspended in phosphate-buffered saline (PBS; Corning, Corning, NY, cat # 21040CV), and gently transferred to tubes containing 10 ml of a 100% polysucrose/sodium diatrizoate solution (Histopaque-1077; Sigma-Aldrich, cat # H8889) overlaid with 15 ml of 75% Histopaque-1077. After centrifugation at 40°C for 30 min at 1200×*g*, the tumor cells in the top interface above the 75% Histopaque were collected and plated on cell culture plates. After establishing cells that grew in a monolayer, the morphology of the cell lines was evaluated by microscopy.

UPS cells were cultured in DMEM supplemented with 20% FBS, 1% penicillin/streptomycin (100 U/ml), and 1% L-alanyl-L-glutamine (glutagro; Corning, cat # 25-015-CI). The UPS060.1 and UPS271.1 nomenclature denotes that these cell lines were established from a patient-derived xenograft tumor that arose from a subcutaneous injection of either UPS060 cells or UPS271 cells into a mouse. MPNST007 cells were cultured in DMEM/F12 (Corning, cat # 10090CV) supplemented with 15% FBS, 1% penicillin/streptomycin (100 U/ml), 1% glutagro, and 1% nonessential amino acids (Corning, cat # 25-025-CI). MPNST3813E cells were cultured in DMEM supplemented with 10% FBS and 1% penicillin/streptomycin (100 U/ml). Isolation of MPNST642 cells was described previously^38^, and this cell line was characterized further in this study by sequencing analysis. Cells were cultured in a humidified incubator at 37°C with 5% CO_2_. Cells were tested for mycoplasma every 6 months.

Doubling time was measured by determining cell confluence using an Incucyte system (Essen Bioscience, Ann Arbor, MI). UPS or MPNST cells were plated in 24-well plates or 96-well plates at 10% to 30% confluence and imaged every 4 h until they reached confluency. The doubling time was calculated using a nonlinear regression fit for exponential growth in Prism 7 software (GraphPad Software, La Jolla, CA).

Short tandem repeat (STR) DNA profiling was performed using a Powerplex 16 HS kit (Promega, cat # DC2101). Genomic DNA was extracted from the tumor specimens and the cultured cell lines using a QIAmp mini DNA isolation kit (Qiagen, cat # 51304). Sixteen loci were screened for regions of microsatellite instability with defined trinucleotide or tetranucleotide repeats within the chromosomes. The STR results for each cell line were authenticated against the corresponding primary tumor’s STR profile and internal database of over 4000 public profiles and profiles unique to cell lines developed by MDACC investigators. To comply with the Health Insurance Portability and Accountability Act of 1996 and MD Anderson institutional policy, only a selection of human loci is shown. The alleles from 8 human loci are reported in this study: AMEL, CSF1PO, D13S317, D16S539, D5S818, D7S820, TH01, and TPOX.

Normal Schwann cells (NSCs), considered the cell of origin for MPNST, served as control cells. NSCs were cultured in Schwann Cell Medium (ScienCell Research Laboratories, cat # 1700 and 1701). Human bone-marrow-derived mesenchymal stem cells were cultured in mesenchymal stem cell medium (PromoCell cat # C12974 and C28009) and used as a reference for *MDM2* amplification, as they are a potential cell of origin for UPS. DDLPS-224A cells were cultured in DMEM supplemented with 20% FBS and 1% penicillin/streptomycin (100 U/ml) and used as a positive control for *MDM2* amplification.^48^

### Clonogenicity and migration assays

Clonogenicity was assayed as described by May et al. ^21^ Briefly, cells were seeded onto 6-well plates at low densities of 5000 cells for UPS cell lines and 500 to 2,000 cells for MPNST cells (depending on the cell line) and were allowed to form colonies for 14 to 30 days. Images were obtained using a digital scanner (Konica Minolta, Tokyo, Japan) or a BZ-X800 microscope (Keyence Corporation, Itasca, Illinois). Migration assays were performed using a modified Boyden chamber assay with Transwell inserts containing polyethylene terephthalate membranes with 8-μm pores (Corning, cat # 354578). For clonogenicity and migration assays, cells were fixed with 5% glutaraldehyde and stained with 0.1% crystal violet in 20% methanol. For migration assays, Images at 20× magnification were taken with a Nikon microscope (Tokyo, Japan) or a BZ-X800 microscope (Keyence Corporation, Itasca, Illinois).

### Protein expression assays

Sodium dodecyl sulfate-polyacrylamide gel electrophoresis and Western blotting were performed as described by May et al. ^21^ Briefly, proteins were extracted from cultured cells in passive lysis buffer (Promega, cat # E1941) containing protease inhibitors (Sigma-Aldrich, cat # 11697498001) and phosphatase inhibitors (Sigma-Aldrich, cat # P5726-1ML, P0044-1ML). Thirty micrograms of protein was loaded on to precast polyacrylamide gels (Bio-Rad, cat# 4561085, 4568023), 4% to 15% precast polyacrylamide gels for PRC2 components and p53. Acid extraction was performed on cultured cells for isolation of histones, and 3 μg of protein was separated on 4% to 20% precast polyacrylamide gels (Bio-Rad, cat # 4561095) for detection of histone H3 andH3K27me3. Proteins were transferred onto low-fluorescence polyvinylidene fluoride membranes using a semidry transfer method according to the manufacturer’s instructions (Bio-Rad, cat # 1704274). Membranes were blocked in 5% nonfat dry milk before probing with the specific antibodies listed in Supplemental Table 1. Horseradish peroxidase-conjugated secondary antibodies were detected by enhanced chemiluminescence (GE Healthcare, cat # RPN2236) and imaged using a Chemidoc imager (Bio-Rad).

### *In vivo* xenograft studies

The MD Anderson Institutional Animal Care and Use Committee approved all animal procedures. We proceeded with patient-derived xenograft (PDX) implant generation when we received < 1 gram of residual tumor; if the residual tumor weighed > 2 grams, we generated a PDX implant and established a cell line. For PDX implantation experiments, 6-week old male or female NOD.Cg-*Prkdc^scid^ Il2rg^tm1Wjl^*/SzJ (NSG) mice (Jackson Laboratory, Bar Harbor, ME) were implanted with pieces of MPNST007, MPNST3813E, RIS620, or UPS0103 tumor (approximately 3-5 mm in diameter) in the flank. For injections of UPS060.1 cells, 3 × 10^6^ cells in 1:1 PBS:Matrigel (Corning, cat #354248) (0.1 ml/mouse) Trypan blue viable cells were injected subcutaneously into the flank of 6-week old male or female NSG mice. For the UPS511 cell line, 5 × 10^6^ Trypan blue viable cells in 1:1 PBS:Matrigel (Corning, cat #354248) (0.1 ml/mouse) were injected subcutaneously into the flank of 6-week old male or female NSG mice. Mice were monitored for tumor size, body weight, and well-being and were euthanized when tumors reached an approximate volume of 1000-2000 mm^3^. Tumor growth was measured once per week using a caliper. The mice were not assessed for metastases. Figures show data from at least 4 representative animals. Resected tumors were weighed, fixed in formalin and paraffin-embedded for IHC studies. IHC studies including hematoxylin and eosin were performed to compare the xenograft specimens to the original patient tumors. The percentage uptake was calculated by dividing the number of mice that grew tumors by the total number of mice implanted or injected with cells. The latency period was estimated from the tumor growth data.

### Sequencing

Targeted sequencing was performed for a panel of 7 genes frequently mutated in UPS and MPNST according to TCGA: *TP53, NF1, ATRX, CDKN2A, EZH2, EED*, and *SUZ12*, using genomic DNA isolated from cultured cells with a Qiagen QIAmp mini DNA isolation kit ^28, 49, 50^. DNA was fragmented and bait-captured according to the manufacturer’s guidelines ^51^. Libraries were sequenced on a HiSeq2000 or HiSeq4000 sequencer (Illumina, San Diego, CA) with 76-bp paired-end reads for MPNST007, MPNST3813E, UPS0103, and RIS620; and 76-bp single reads for UPS060.1, UPS271.1, and UPS511. The raw data were converted to fastq format and aligned to the reference genome (hg19) with the Burroughs-Wheeler Aligner ^52^. Median target coverage was 256× to 763× for MPNST007, MPNST3813E, MPNST4970; and UPS0103 cells and 207× to 230× for UPS060.1, UPS271.1, and UPS511 cells. Mutations were called against a common normal lung sample using the MuTect and Pindel tools ^53, 54^. Sanger sequencing of *TP53* was performed for genomic DNA isolated from the original MPNST specimen using either an ABI 3730XL or 3730 DNA sequencer (Thermo Fisher Scientific, Waltham, MA) with a Big Dye Terminator cycle sequencing chemistry kit (Thermo Fisher Scientific, cat # 4337455) according to the manufacturer’s guidelines.

### Copy number alteration analysis

*MDM2* copy number analysis was performed using the MDM2 TaqMan Copy Number Assay Hs02970282_cn (Thermo Fisher Scientific cat # 4400291) and the reference assays for TERT (Thermo Fisher Scientific cat # 4403316) and RNase P (Thermo Fisher Scientific cat # 4403326) according to manufacturer’s guidelines. Results were analyzed using CopyCaller Software v2.1 (Thermo Fisher Scientific, Waltham, MA).

## RESULTS

### Establishment and morphology of UPS and MPNST cell lines

All cell lines were established from cells isolated from residual resected tumors. The UPS cell lines were generated from recurrent or metastatic tumors, and the MPNST tumors (all 3 NF1-associated) were generated from primary tumors (Table 1). A sarcoma pathologist at MD Anderson Cancer Center confirmed the final histology of the tumors. The STR profiles of the established cell lines matched those of the original tumors, indicating that each of the cell lines was derived from the tumor of origin, and there was no cross-contamination with other cell lines. The hematoxylin and eosin staining of the original specimens showed a mixture of tumor cells and stromal cells in UPS and MPNST tumors (Figure 1a, lower panels). The cells were all mononuclear. UPS060.1 and UPS511 cell lines had an elongated spindle-like shape, while UPS271.1 and UPS0103 cell lines were polygonal and RIS620 cells exhibited both polygonal and spindle shapes (Figure 1a, upper panels). MPNST007 cell lines had an elongated spindle shape, whereas MPNST3813E cells showed more polygonal morphology in comparison, as did MPNST4970 cells (Figure 1a, upper panels).

**Table 1.** Characteristics of UPS and MPNST patients from whom tumors were resected and cell lines derived.

| **Cell Line** | **Date of Surgery** | **Sex** | **Age (y)** | **Race/ethnicity** | **Tumor Location** | **Final Histology** | **Status** |
| --- | --- | --- | --- | --- | --- | --- | --- |
| RIS620 | 2011 | F | 50 | W | Right chest wall and scapula | RAD-UPS | R |
| UPS060 | 2010 | F | 8 | B | Pelvis, involving preserosal soft tissue of colon, bladder wall, and right ureter | UPS | R |
| UPS271 | 2010 | M | 60 | H | Right anterior thigh | UPS | R |
| UPS511 | 2011 | M | 73 | W | Chest wall | UPS | R |
| UPS0103 | 2016 | M | 37 | W | Paraspinal, L1 | UPS | M |
| MPNST007 | 2011 | F | 50 | B | Left calf | MPNST | P |
| MPNST3813E | 2016 | F | 20 | H | Left buttock | MPNST | P |
| MPNST4970 | 2018 | M | 28 | W | Foot | MPNST | R |

Abbreviations: W, white; B, black; H, Hispanic; RAD, radiation-associated; UPS, undifferentiated pleomorphic sarcoma; MPNST, malignant peripheral nerve sheath tumor; R, recurrence; M, metastasis; P, primary tumor

**Table 2.** Short tandem repeat (STR) fingerprinting of UPS and MPNST cell lines.

| Tumor or cell line | STR loci* |  |  |  |  |  |  |  |
| --- | --- | --- | --- | --- | --- | --- | --- | --- |
|  | AMEL | CSF1PO | D13S317 | D16S539 | D5S818 | D7S820 | TH01 | TPOX |
| UPS060 tumor | X | 11,12 | 10,11 | 11,13 | 10,12 | 9 | 8,9 | 9,12,13 |
| UPS060 cell line | X | 11,12 | 10,11 | 11,13 | 10,12 | 9 | 8,9 | 9,13 |
| UPS0103 tumor | X,Y | 9,10 | 9,11 | 9,10 | 12 | 8,10 | 6,7 | 8,11 |
| UPS0103 cell line | X,Y | 9,10 | 9,11 | 9,10 | 12 | 8,10 | 6,7 | 8,11 |
| MPNST007 tumor | X | 8,10 | 11 | 10,13 | 11,12 | 11,12 | 7,9.3 | 9,11 |
| MPNST007 cell line | X | 8,10 | 11 | 10,13 | 11,12 | 11,12 | 7,9.3 | 9,11 |
| MPNST3813E tumor | X | 12 | 8,12 | 13 | 11 | 8,11 | 9 | 9,11 |
| MPNST3813E cell line | X | 12 | 8,12 | 13 | 11 | 8,11 | 9 | 9,11 |
| MPNST4970 tumor | X | 11 | 11 | 14 | 12 | 10 | 9.3 | 8 |
| MPNST4970 cell line | X | 11 | 11 | 14 | 12 | 10 | 9.3 | 8 |

\*To comply with the Health Insurance Portability and Accountability Act of 1996 and MD
Anderson institutional policy, only a selection of markers is shown. Marker selection was based on ATCC's STR database. STR results for UPS271, UPS511, and RIS620 are available in May et al. Cancer Biology and Therapy, 2017.

**Figure 1.**
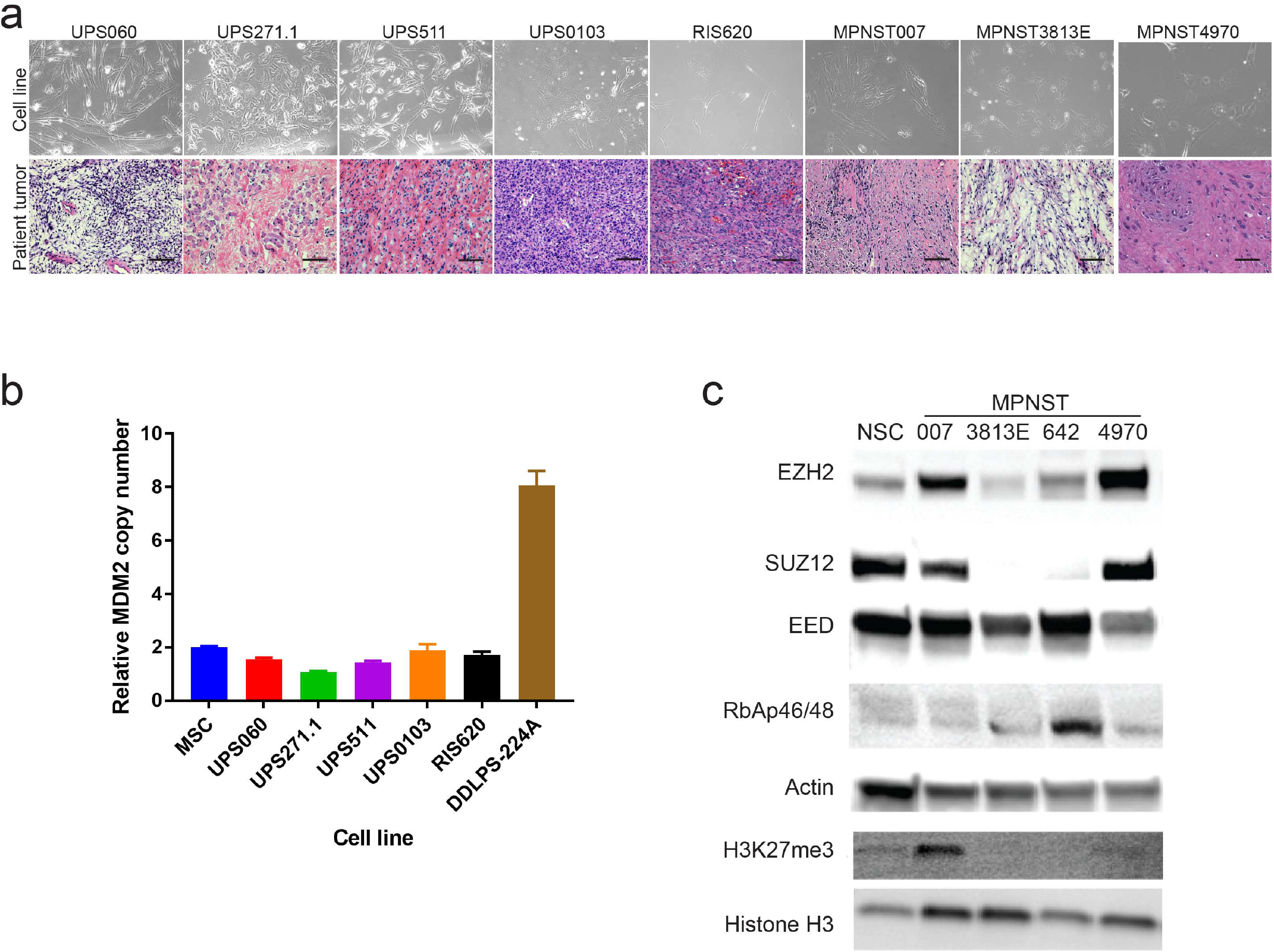
Characterization of cell lines and original tumors. (a) Bright-field images of UPS and MPNST cell lines (upper panels) and images of hematoxylin and eosin (H&E)-stained sections of the original patient tumors (lower panels), 20× magnification. Scale bar represents 100 μm. (b) *MDM2* copy number analysis in UPS cell lines. MSC, mesenchymal stem cells, were used as a reference. DDLPS-224A cells were used as a positive control for *MDM2* amplification. *TERT* was used as the reference gene for the *MDM2* copy number assay. N = 2, graph shows calculated copy number; error bars indicate copy number range. (c) Immunoblots showing expression of EZH2, SUZ12, EED, RbAP46/48, and H3K27me3 in normal Schwann cells (NSC), MPNST007, MPNST3813E, MPNST642, and MPNST4970 cells. EZH2, SUZ12, EED, and RbAP46/48 levels were normalized to that of actin (middle panels). H3K27me3 expression was normalized to H3 levels (lower two panels).

As of May 2021, UPS060.1 has been carried to passage 50, UPS271.1 to passage 36, UPS511 to passage 35, and UPS0103 to passage 17. MPNST007 and MPNST3813E have been taken over passage 100, MPNST4970 has been carried to passage 41. Our laboratory will continue to passage the lower passage UPS and MPNST cell lines to determine whether they can be carried to over 90 passages.

The possibility of misclassification of the UPS tumors as dedifferentiated liposarcoma (DDLPS) was ruled out by *MDM2* copy number analysis using genomic DNA isolated from the established cell lines. Bone marrow-derived mesenchymal stem cells served as the reference for *MDM2* copy number, as they represented a potential cell of origin for UPS. DDLPS-224A, a DDLPS cell line with known amplification of *MDM2*, served as a positive control ^48^. No amplification of MDM2 was detected in the UPS cell lines, providing evidence that the specimens from which these cell lines were derived were indeed UPS and not missclassified DDLPS.

### Protein expression

Western blotting was used to assess the expression of histone marks and proteins known to be associated with MPNST: H3K27me3, and the PRC2 members SUZ12, EED, RbAp46/48, and EZH2. Normal Schwann cells (NSCs), considered the cell of origin for MPNST, served as control cells. The MPNST007 and MPNST4970 cell lines expressed higher levels of EZH2 than did the control NSCs. The MPNST cells were also assayed for the absence of H3K27me3, which is considered a potential diagnostic marker of NF1-associated MPNSTs ^20^. The H3K27me3 mark was present in MPNST007 cells and MPNST4970 but not in MPNST642 or MPNST3813E cells (Figure 1c, lower panels). These findings suggested that MPNST007 and MPNST4970 are PRC2-competent and MPNST642 and MPNST3813E are PRC2-incompetent.

### Cell proliferation rate

The growth rate for the UPS and MPSNT cell lines was determined by measuring relative confluence (Figure 2a-b). For the UPS cell lines, the doubling time ranged from approximately 33.3 to 279.8 h. UPS060.1 cells had the shortest doubling time (33.3 - 53.5 h), and the doubling time of UPS271.1 cells was similar to that of UPS060 cells (48.8 - 56.6 h). UPS511 cells and RIS620 cells had intermediate doubling times (90.6 - 92.0 h and 151.5-174.8 h, respectively). The UPS0103 cell line had the slowest doubling time of the UPS cells in this study, at 279.8 h. Of the MPNST cell lines, MPNST3813E cells had a doubling time of 36.3 h (rangerange 34.6 to 39.7 h). MPNST007 cells doubled in 49.4 to 58.9 h, and MPNST4970 doubled in approximately 84.6 h. These results showed that the cell lines established in the current study have a wide range of doubling times, which reflects the heterogeneous nature of these tumors.

**Figure 2.**
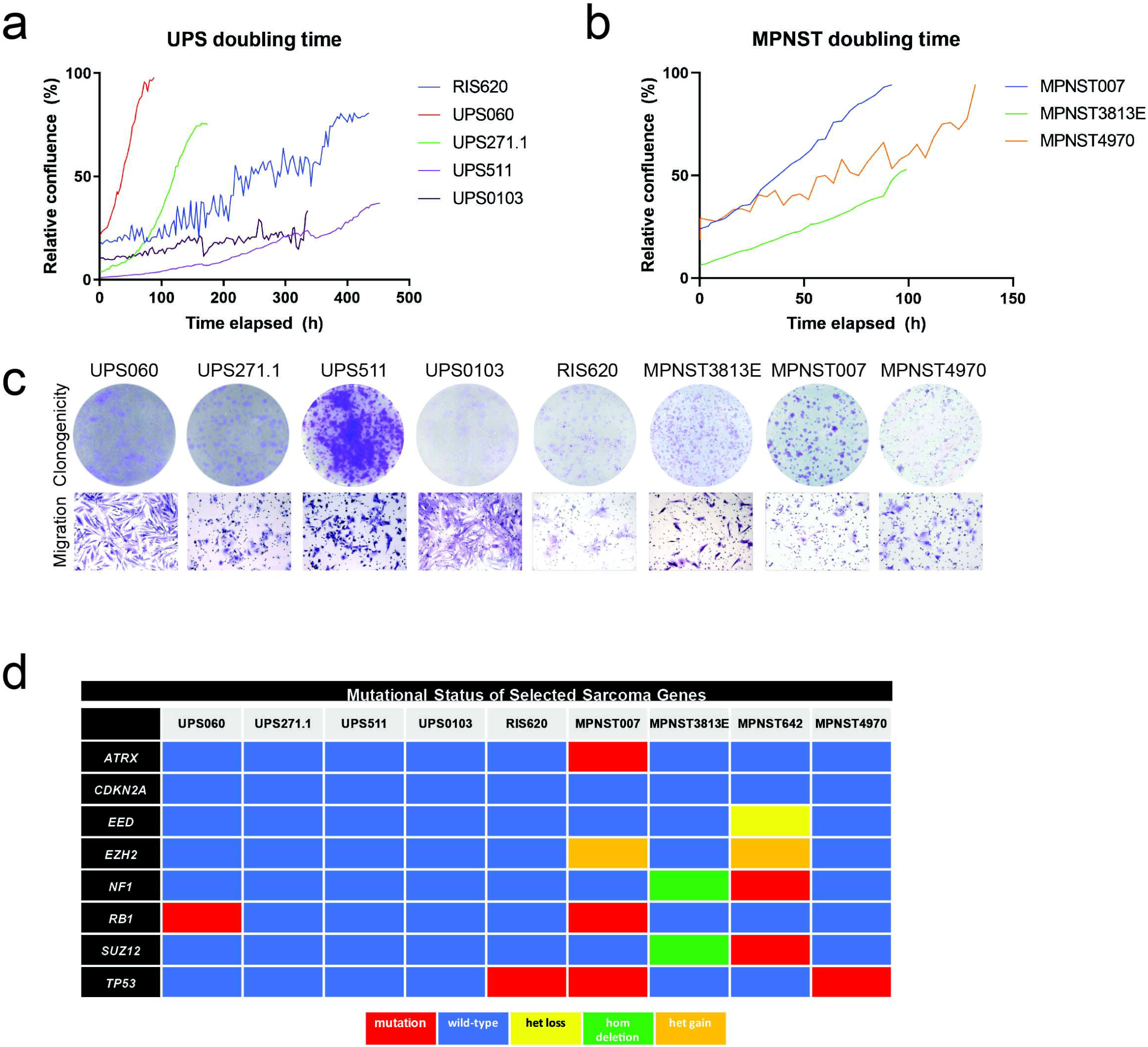
*In vitro* growth studies of UPS and MPNST cells. Doubling time for UPS (a) and MPNST (b) cell lines. Cells were plated and allowed to grow until they reached a growth plateau. Relative confluence was measured using an Incucyte device (Essen Bioscience). (c) Clonogenicity assay for UPS and MPNST cells. For UPS cells, 5000 cells were seeded and allowed to grow for the indicated number of days. For MPNST cells, 500 to 2000 cells were seeded and allowed to grow for 16 days. Migration of UPS cells and MPNST cells, 20× magnification. (d) Targeted sequencing results for selected sarcoma genes. Refer to the results section for details on the mutations identified.

### *In vitro* tumorigenic potential of the new UPS and MPNST cell lines

The UPS and MPNST cell lines were assessed for clonogenic and migratory potential. For clonogenic assays, cells were seeded at a low density and allowed to form colonies for up to 16 days. All of the UPS cell lines formed colonies. UPS060, UPS271.1, and RIS620 formed small colonies, UPS511 formed large distinct colonies, and UPS0103 formed diffuse colonies (Figure 2c). MPNST007, and MPNST3813Eformed distinct colonies, while MPNST4970 cells formed more diffuse colonies. Further analysis of tumorigenic potential was performed by measuring the migration of the established cell lines in a modified Boyden chamber assay. All the UPS and MPNST cell lines exhibited the ability to migrate toward a chemoattractant (Figure 2d, upper panels). These results showed that the newly established cell lines had the ability to form colonies, and migrate.

### DNA mutation analysis of selected cancer genes

The mutation status of genes frequently mutated in UPS and MPNST according to the TCGA, including *ATRX, CDKN2A, EED, EZH2, NF1, RB1, SUZ12*, and *TP53*, was assessed using high-coverage targeted sequencing with a minimum median target coverage of approximately 200× ^28, 49, 50 55^. Figure 2d summarizes the mutations found in the cell lines. UPS271.1, UPS511, UPS0103 cell lines had no mutations in the cancer genes listed above. In the UPS060 cell line, a mutation in the *RB1* gene was identified: c.C1574G; p.A525G. This mutation has been reported in hemangioblastomas and acute myeloid leukemia ^56, 57^. The RIS620 cell line carried a *TP53* mutation at c.G818A; p.R273H, that was documented in lung cancer, esophageal cancer, pancreatic carcinoma, hematological malignancies, ovarian and endometrial cancers, and gastric cancer ^23, 58–63^. The MPNST007 cell line had a mutation in *ATRX* at c.6338_6341del; p.F2113fs; this mutation was previously found in neuroblastoma, pancreatic neuroendocrine tumors, and uterine leiomyosarcomas ^64–66^. The MPNST007 cell line also carried a mutation in *RB1*: c.71_72del; p.P24fs, which to our knowledge has not been reported previously. A truncating mutation in *TP53*, c.301A>T; p.K101*, was identified in the MPNST007 original tumor, but this mutation was not found in the MPNST007 cell line sequencing results. Instead, a different *TP53* mutation was identified: c.965delC; p.322fs, which was also reported in colon and rectal cancer and prostate cancer ^67, 68^. MPNST3813E cells had copy number loss of *NF1* and *SUZ12*, and MPNST4970 cells had *TP53* stopgain at c.C97T:p.Q33X. The MPNST642 cell line carried previously unreported mutations in *NF1* at c.511delA; p.N171fs and in *SUZ12* at c.2026dupA; p.D675fs.

### *In vivo* tumorigenicity studies

The *in vivo* tumor-forming capacity of the established cell lines was determined by either subcutaneous injection or tumor implantation. For UPS060.1, and UPS511 cell lines, 3-5 million cells were injected subcutaneously into the flank of NSG mice, and tumor growth was measured weekly (Figure 3a-b). For the implantation experiments, UPS0103, RIS620, MPNST007, and MPNST3813E, tumor pieces of approximately 3-5-mm^3^ were embedded into the flank of NSG mice, which were then monitored for tumor growth (Figure 3c-f). Mice were sacrificed when the tumors reached a volume of 1000 to 2000 mm^3^. Hematoxylin and eosin staining of the xenograft tumors was performed on slides cut from formalin-fixed, paraffin-embedded blocks (Figure 3a-g, right panels). Cells from the xenografts had similar morphology as cells from the parental tumors. The percentage of uptake of tumor cells and latency period are shown in Table 3. Overall, the cell line injection and implant xenograft models had uptake ranging from 47% to 100%. The latency period was the longest for MPNST3813E cells (115 days) cells and the shortest for RIS620 cells (23 days) (Table 3). The UPS 271.1 and MPNST4970 cell lines show a heterologous growth pattern for generation of xenografts that is currently under investigation.

**Figure 3.**
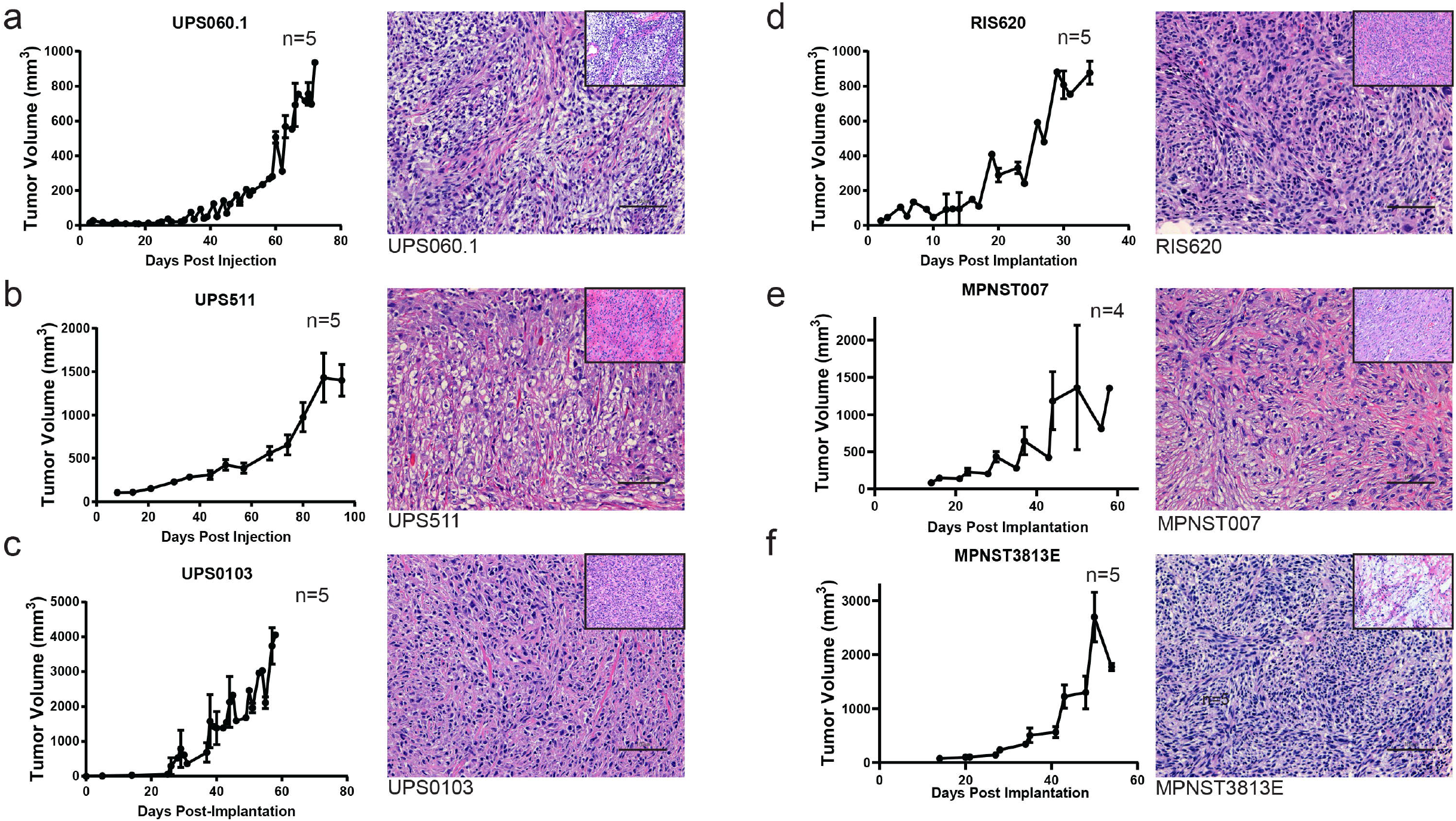
*In vivo* xenograft models of UPS and MPNST. (a-d) Tumor growth after UPS cell line injection or implantation in 5 representative animals (left panels). Images of hematoxylin and eosin (H&E)-stained sections of xenograft tumors at 20× magnification (right panels). Scale bar indicates 100 μm. Inset shows original patient tumor image at 20× magnification. (e-g) Tumor growth after MPNST cell line implantation in 5 representative animals (left panels). Images of H&E-stained sections of xenograft tumors at 20× magnification (right panels). Scale bar indicates 100 μm. Inset shows original patient tumor image at 20× magnification.

**Table 3.** Latency and uptake for UPS and MPNST xenograft models.

| Cell line/implant | % uptake | Latency (days)<sup>a</sup> |
| --- | --- | --- |
| UPS060.1 | 70 | 37 |
| UPS511 | 100 | 50 |
| UPS0103 | 77 | 30 |
| RIS620 | 72 | 23 |
| MPNST007 | 47 | 46 |
| MPNST3813E | 70 | 32 |
| MPNST4970 | 90 | 28 |
| MPNST4970 SQL | 50 | 146 |

<sup>a</sup> Determined from Figure 3a-g tumor growth graphs.

## DISCUSSION

Primary tumor cell lines established from patient tumors can recapitulate the original patient tumor in an *in vitro* setting to investigate molecular targets further. The goal of the present study was to generate and characterize cellular and xenograft models to study of 2 genomically complex sarcomas, UPS and MPNST. We successfully established 5 UPS cell lines. UPS was previously classified as malignant fibrous histiocytoma (MFH); the disadvantage of using MFH cell lines for the study of UPS is that the now-defunct term MFH encompassed tumors with 4 different histologies—myxofibrosarcoma, pleomorphic liposarcoma, rhabdomyosarcoma, and UPS ^1^. Therefore, researchers require cell lines with pathological confirmation to represent UPS. The UPS cell lines described in this study were derived from patient tumors classified as UPS by a sarcoma pathologist at MD Anderson Cancer Center.

STR profiling showed each cell line matched the STR profiles of the parental tumors with no signs of contamination with other cell lines. The establishment of cancer cell lines presents some difficulties; most primary cultures are unsuccessful because cells senesce after several months in culture. A recent study had a moderate success rate of 15% in establishing sarcoma cell lines from resected tumors ^37^. The soft tissue sarcoma cell lines described here have been in culture for approximately 1-2 years at this writing. The cell lines all underwent spontaneous transformation and were not immortalized by the introduction of large T-cell SV40 antigen or human telomerase reverse transcriptase. Moreover, the established cell lines successfully formed xenograft tumors when administered to immunocompromised mice.

Targeted sequencing was performed on selected known cancer-related genes to identify genetic alterations in the established UPS and MPNST cell lines and provide an initial genomic characterization of the cell lines. Our finding that the genomes of the UPS and MPNST cell lines were not highly mutated agreed with the recent TCGA study of sarcomas ^28^. In that study, the authors reported copy number losses in *ATRX, CDKN2A, NF1, RB1*, and *TP53*. Interestingly, we found no mutations in *NF1* in MPNST007, MPNST3813E, MPNST4970 cell lines. However, these cell lines may have copy number alterations of *NF1* that cannot be inferred from the targeted sequencing analyses. PRC2 mutations, specifically inactivation of *EED* and *SUZ12*, have been documented in MPNST ^69–71^. Of the 3 NF1-associated MPNST cell lines characterized in this study, MPNST642 carried a frameshift mutation (p.D675fs) in *SUZ12*. The absence of *EZH2* mutations in the NF1-MPNST cell lines described here, supports the finding that *EZH2* function is needed for uncontrolled tumor growth and that its expression is significantly upregulated in MPNST specimens compared to normal nerve cells ^72, 73^. Cell lines containing functional PRC2 and cell lines with inactive PRC2 are needed to elucidate the contribution of functional PRC2 to MPNST progression, as the full relevance of this complex in a therapeutic setting requires further study. In the current study, we chose not to evaluate the cell lines’ sensitivity to commonly used chemotherapeutic agents as MPNST patients do not respond; thus, there is a low chance the new cell lines will be sensitive to treatment.

In summary, we successfully established several new cellular models for the study of UPS and MPNST. Furthermore, the initial characterization of the cell lines showed that they could be used as *in vitro* and *in vivo* models for further study of sarcoma etiology. Analysis of mutations in genes known UPS and MPNST genes showed that the cell lines recapitulate known genetic alterations in UPS and MPNST. We hope that the establishment of these cellular models will allow researchers to take the next steps in investigating these debilitating soft tissue sarcomas in greater detail.

## Acknowledgements

The authors acknowledge the Transgenic Mouse Facility and the Surgical Oncology Histology Facility at MD Anderson Cancer Center. The authors also thank Amy Ninetto and the Department of Scientific Publications at MD Anderson Cancer Center for their assistance in editing this manuscript.

## Author Contributions

Author contributions: Angela D. Bhalla: conceptualization, data interpretation, experiments, Sharon M. Landers: cell line generation, experiments, Anand K. Singh: experiments, Michelle G. Yeagley: cell line generation, chart review, patient cconsenting, experiments, Gabryella S.B. Serrrao: cell line generation, chart review, patient consenting, Zachary A. Mulder: chart review, patient consenting, Jace P. Landry: experiments, chart review, Cristian B. Delgado-Baez: experiments, Stephanie Dunnand: experiments, Veena Kochat: experiments, Katarzyna J. Tomczak: data mining, Theresa Nguyen: cell line generation, experiments, XiaoYan Ma: experiments, Svetlana Bolshakov: experiments, Brian A. Menegez: cell line generation, experiments, Salah E. Lamhamedi Cherradi: cell line generation, experiments, Joseph A. Ludwig: conceptualization, Hannah C. Beird: data mining, Xizeng Mao: bioinformatics analysis, Xingzhi Song: bioinformatics analysis, Davis R. Ingram: experiments, chart review, patient consenting, Wei-Lien Wang: pathological analysis, Alexander J. Lazar: pathological analysis, Ian E. McCutcheon: chart review, patient care, John M. Slopis: chart review, patient care, Kunal Rai: conceptualization, Jianhua Zhang: conceptualization, Dina Lev: conceptualization, Keila E. Torres: conceptualization, data interpretation, funding acquisition, project administration.

## Data Availability Statement

All data collected and analyzed in this study are available.

