## Supplemental Figure 1 and Tables 1-4 for "New cellular models of undifferentiated pleomorphic sarcoma and malignant peripheral nerve sheath tumor"

**SUPPLEMENTARY INFORMATION**

Supplemental methods

Quantitative Real-Time PCR (qPCR). PCR amplifications were performed on the Light Cycler 480 II (Roche, Indianapolis, IN, USA). A 20uL reaction containing 50ng of DNA was amplified in duplicate with either *MDM2* primers (Accession number NG_016708.1, Forward: 5'-AAAGGGCCAGGTTAAATGGT-3', and Reverse: 5'-GTGTGCCCCAGAACAAAGAT-3') or *GAPDH* primers (Accession number NC_018923.2, Forward 5'-AAGACAGAATGGAAGAAATGTGC-3', and Reverse 5'-GAGATGGGGACAGGACCATA-3') and the Light Cycler 480 SyberGreen master mix. The cycling profile was as follows: pre-incubation at 50°C for 2 minutes, denaturation at 95°C for 5 minutes, 45 cycles of amplification , 95°C 15 seconds, 60°C for 30 seconds, melting curve. Quantification Cycle (Cq) was calculated by the instrument using the absolute quantification/second derivative. Relative copy number was determined by 2−ΔΔCT using human bone marrow-derived mesenchymal stem cells as a control and *GAPDH* as the reference gene. The cut off for MDM2 amplification was 5.0.

Immunohistochemical studies. Slides of the formalin-fixed paraffin-embedded tissues were stained for pankeratin, S-100, desmin, and smooth muscle actin (SMA) by the Research Histology Clinical Laboratory at MD Anderson Cancer Center which is supported by National Institutes of Health through Cancer Center Support Grant (CCSG) P30 CA016672.

Supplemental Figure 1.

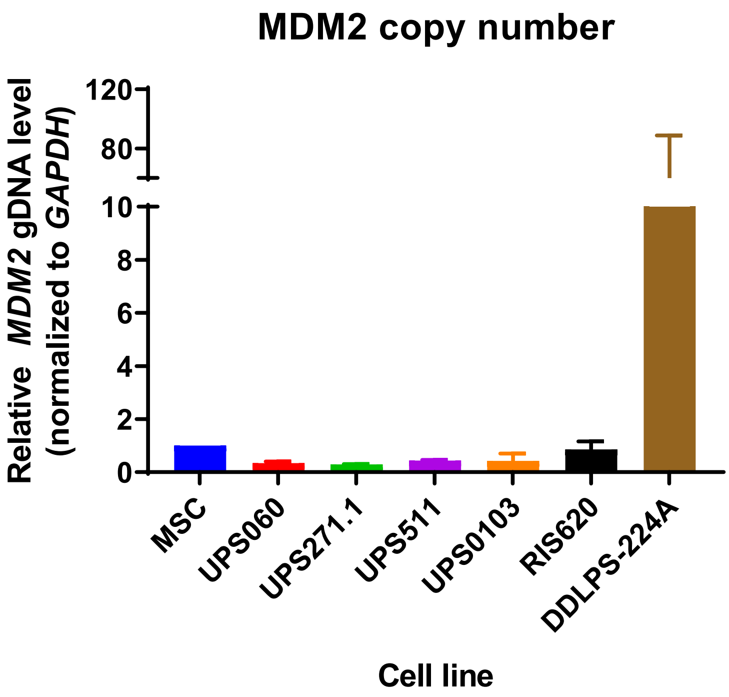


Quantitative real-time rt PCR for MDM2 copy number using gDNA isolated from UPS cell lines.

N=2, GAPDH was used as control for copy number.

Supplemental Table 1. Antibodies used in this study.

| **Antibody** | **Vendor and catalog number** | **Dilution** |
|---|---|---|
| anti-SUZ12 | Cell Signaling Technologies, 3737S | 1:1000 |
| anti-EZH2 | Cell Signaling Technologies, 5246S | 1:1000 |
| anti-EED | Sigma-Aldrich, 09-774 | 1:500 |
| anti-RbAp46/48 | Cell Signaling Technology, 4688S | 1:1000 |
| anti-H3 | Cell Signaling Technologies, 4499S | 1:1000 |
| anti-H3K27me3 | Cell Signaling Technologies, 9733S | 1:1000 |
| anti-rabbit HRP-conjugated secondary | Santa Cruz, sc-47778<br>GE, NA9341ML | 1:3000<br>1:10000 |

Abbreviation: HRP, horseradish peroxidase

Supplemental Table 2. Pathology notes for UPS patients

| Cell Line | Primary Tumor Pathology Report | Comments (on later pathology reports) |
|---|---|---|
| **RIS620** | UNCLASSIFIED PLEOMORPHIC HIGH-GRADE SPINDLE CELL SARCOMA (no comments on tests) | No molecular testing identified in any path reports with sarcoma diagnosis |
| **UPS060** | Immunohistochemical studies performed at UTMDACC reveal that the tumor is negative for pankeratin, S-100, desmin, SMA, myogenin, and MyoD1. These findings support a mesenchymal origin are not specific to a type. These findings do not support a rhabdomyosarcoma. Overall, this tumor is best categorized as an unclassified spindle cell sarcoma and is favored to be high-grade. Per outside report, the tumor cells were reported to be strongly positive for vimentin, with focally positivity for CD34, laminin and type IV collage, while being negative for desmin, myogenin, caldesmon, CAM 5.2, and S-100. | |
| **UPS271** | The tumor has a multinodular growth pattern and appears plexiform in areas. Vascular invasion is notable. Immunostains prepared here for keratin, S-100, and myogenin are negative. Desmin is positive, but this is regarded as a nonspecific finding. | |
| **UPS511** | Submitted immunohistochemical stains show that the tumor is positive for vimentin and focally positive for CD34, SMA and CD68. It is negative for CD117, S-100 protein, desmin, keratin and TTF-1. Ki-67 positive in approximately 25% of tumor cells. These findings are compatible with a myxofibrosarcoma. | |
| **UPS0103** | The performed immunohistochemical stains show tumor cells are negative for pankeratin, S100 protein, SMA, desmin, CD31, CD34 and ERG. The above diagnosis is unchanged. | |

Supplemental Table 3. Clinical characteristics of MPNST patients.

| Cell Line | Clinical Characteristics |
|---|---|
| **MPNST007** | multiple plexiform and cutaneous neurofibromas, scoliosis, and two café au lait spots |
| **MPNST3813E** | café au lait spots, family history of NF1, plexiform neurofibroma, minor learning disabilities, seizures, scoliosis, and leg length discrepancy |
| **MPNST642** | café au lait spots, ADHD, cutaneous neurofibromas, axillary and inguinal freckling, and plexiform neurofibroma |
| **MPNST4970** | Immunohistochemical studies show low heterogeneous labeling of high grade sarcoma and atypical neurofibromas with H3K27me3, while SOX10 and p53 are negative in both components |

Supplemental Table 4. Immunohistochemical staining of xenografts and patient tumors for selected UPS and MPNST markers

| Sample | Desmin | SMA | pancytokeratin | S100 |
|---|---|---|---|---|
| **MPNST007** | positive | positive | negative | negative* |
| **MPNST007 xenograft** | negative | positive | negative | negative |
| **MPNST3813E** | focally positive* | nuclear positive | negative | focally positive* |
| **MPNST3813E xenograft** | negative | focally positive | negative | sparse positive cells |
| **RIS620** | positive | positive | negative | positive |
| **RIS620 xenograft** | focally positive | sparse positive cells | negative | negative |
| **UPS0103** | negative* | negative* | negative* | negative* |
| **UPS0103 xenograft** | negative, sparse positive cells | focally positive | negative | negative |
| **UPS511** | negative* | focally positive* | negative* | negative* |
| **UPS511 xenograft** | negative, sparse positive cells | focally positive | positive | negative |
| **UPS060** | negative* | negative* | negative* | negative* |
| **UPS060.1 xenograft** | negative | sparse positive cells | negative | focally positive |
| **MPNST4970 is still under investigation** | | | | |

*information taken from pathology report.
